# Benchmarking generative AI and physics based molecular simulation for sampling conformational heterogeneity in T4 Lysozyme

**DOI:** 10.64898/2026.05.10.724101

**Authors:** Soumendranath Bhakat

## Abstract

Wild-type T4 lysozyme (T4L) is used as a benchmark to evaluate conformational sampling across generative AI, AI-accelerated molecular simulation (AMS), and physics-based enhanced molecular dynamics (EMD). A four-state model: exposed/open, exposed/closed, buried/open, and buried/closed; is defined using physically meaningful collective variables. While generative AI methods (AF-cluster, MSA subsampling of AlphaFold2, ConforFold, AlphaFlow, ESMFlow, ConfRover, BioEmu) largely sample only the exposed/open state, AMS integrating generative ensembles with iterative molecular dynamics, recovering all states and reproducing equilibrium populations similar to EMD and experimental smFRET signatures.

## Introduction

Proteins are molecular machines that govern biological function. They are not static structures but interconvert between multiple transiently populated metastable states, each capable of activating distinct signaling pathways. The populations of these states are modulated by environmental perturbations such as temperature, pH, ligand binding, and mutations. Sampling the transiently populated states is therefore central to understanding how proteins encode function, yet it remains one of the hardest problems in structural biology(1–5).

The development of generative AI algorithms such as AlphaFold has enabled accurate prediction of static protein structures from sequence alone, but these algorithms do not sample transiently populated alternate conformational states. A growing subfield of machine learning for structural biology has aimed to address this by predicting conformational ensembles from sequence alone(1). These approaches fall into two subgroups: MSA-based methods, which perturb evolutionary coupling information in multiple sequence alignments to generate structural diversity, and generative models such as diffusion models or flow-matching frameworks, trained on short MD simulations data, which claim to sample transiently populated alternate states. Despite rapid growth in both subgroups, systematic benchmarking against physically rigorous reference data remains absent, making it difficult to assess whether these methods genuinely capture biologically relevant conformational diversity or simply reproduce structural noise.

Physics-based enhanced sampling methods such as metadynamics(6–9), adaptive sampling(10), and replica exchange(11) provide such a reference. They can sample transiently populated alternate states and their relative populations across a diverse range of biomolecular systems. Sampling efficiency is assessed by how well these methods capture state populations along collective variables (CVs) that discriminate transient states from the ground state(12). However, fair comparisons between physics-based enhanced sampling and generative AI methods are rare due to three key challenges. An ideal benchmark system must have experimentally validated transient states with CVs simple enough to distinguish those states from the ground state. It must have sufficient structural data in the Protein Data Bank so that limited training data cannot be used as a confounding factor. And system size must not unfairly disadvantage generative AI methods, which often struggle with larger and more conformationally complex systems.

Wild-type T4 lysozyme (T4L) satisfies all three criteria. It is small (164 residues), has single-molecule FRET (smFRET) data that directly validates transiently populated alternate states, and has been extensively studied by physics-based enhanced sampling methods. Yet, no prior study has systematically compared generative AI methods, AI-accelerated molecular simulation, and physics-based enhanced sampling on the same system using well characterized CVs. We close this gap here.

## Results

We used two sets of CVs to describe the conformational ensemble of T4L. The first is a two-dimensional CV defined by two C*α* distances: d1 (Ser44–Ile150) and d2 (Glu22–Gln141). The Ser44–Ile150 pair was used in smFRET experiments to probe the opening and closing motion of T4L, with d1 < 2.5 nm defining the closed state and d1 > 2.5 nm defining the open state. The Glu22–Gln141 distance serves as an orthogonal CV capturing hinge domain motion. The second CV is the locking coordinate *p*, developed by *Stock and co-workers(13)*, which quantifies the solvent exposure of Phe4 relative to the hinge helix (see Supplementary Information for full definition). A positive value (*p* > 0) indicates that Phe4 is solvent-exposed, while a negative value (*p* < 0) indicates it is buried within the interdomain interface. Combined with d1, this defines a four-state model: exposed/closed, exposed/open, buried/closed, and buried/open. This four-state classification enables quantitative comparison of state populations across all methods.

We benchmarked three classes of methods. The first class comprises generative AI approaches, divided into two subgroups. The first subgroup consists of MSA-based methods: AF-cluster(14), which clusters MSA subsamples as input to AlphaFold2(15); reduced MSA subsampling with AlphaFold2 (rMSA-AF2)(16); and ConforFold(17), a retrained OpenFold2 model designed to predict alternate conformations. The second subgroup consists of generative models trained on structural or simulation data: AlphaFlow(18), ESMFlow(18), ConfRover(19), and BioEmu(20). The second class is AI-accelerated molecular simulation (AMS), a methodological contribution of this work, in which generative AI ensembles seed multiple rounds of unbiased MD simulations that are combined using a Markov state model (MSM)(21, 22) to recover physically refined state populations (**Figure 1**). The third class is physics-based enhanced sampling repurposed from *Miller and co-workers(23)*, combining 5 µs unbiased MD simulations with metadynamics-seeded unbiased MD simulations (hereafter enhanced molecular dynamics, EMD), which serves as the quantitative reference for all comparisons.

**Figure 1.**
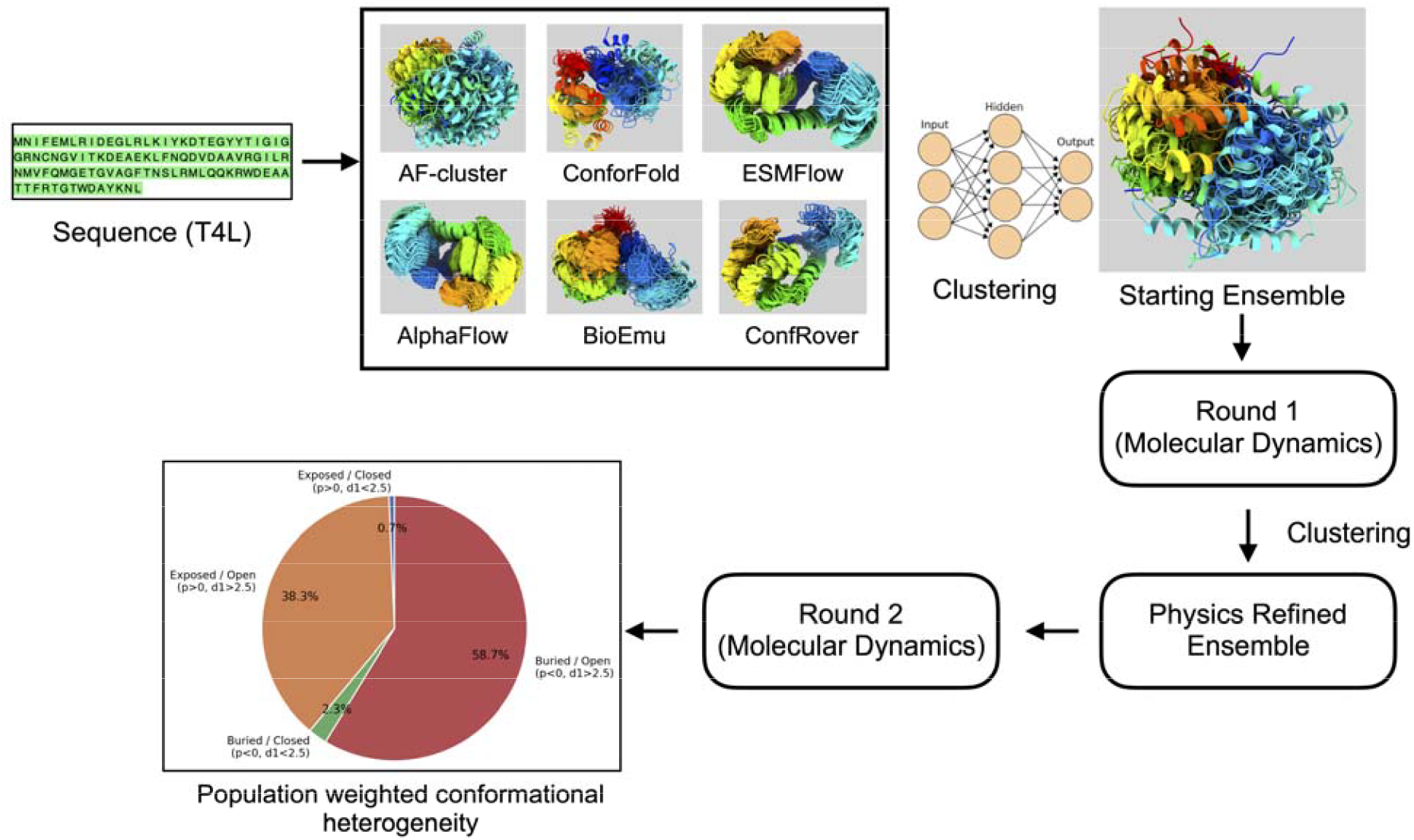
Schematic of the AI-accelerated molecular simulation (AMS) workflow. The workflow begins with the T4L sequence, followed by ensemble generation using multiple generative AI methods, including AF-cluster, ConforFold, BioEmu, ConfRover, ESMFlow, and AlphaFlow. K-center clustering was then performed using transformed distance features to select an initial “starting ensemble” of N=100 structures. Unbiased molecular dynamics simulations of 200 ns were initiated from each member of this starting ensemble in Round 1. The resulting trajectories were clustered using the K-center algorithm on transformed distances, yielding a “physics-refined ensemble” of N=80 structures. Additional unbiased molecular dynamics simulations of 200 ns each were then initiated from the physics-refined ensemble in Round 2. A Markov state model (MSM) was constructed using RMSD features from the combined Round 1 and Round 2 simulations. Equilibrium populations were then projected onto different collective variables to predict conformational populations and heterogeneity in T4L.

AMS successfully sampled all four conformational states: exposed/open, exposed/closed, buried/open, and buried/closed. The two dominant states, exposed/open (AMS: 38.3%, EMD: 39.1%) and buried/open (AMS: 58.7%, EMD: 54.0%), were recovered with relative populations comparable to EMD. The transiently populated buried/closed state was also recovered with a comparable population (AMS: 2.3%, EMD: 2.9%). The exposed/closed state, however, was underrepresented in AMS relative to EMD (AMS: 0.7%, EMD: 3.9%), suggesting that this particular transient state remains the most challenging to access without explicit enhanced sampling to seed follow up unbiased simulations (**Figure 2**).

**Figure 2.**
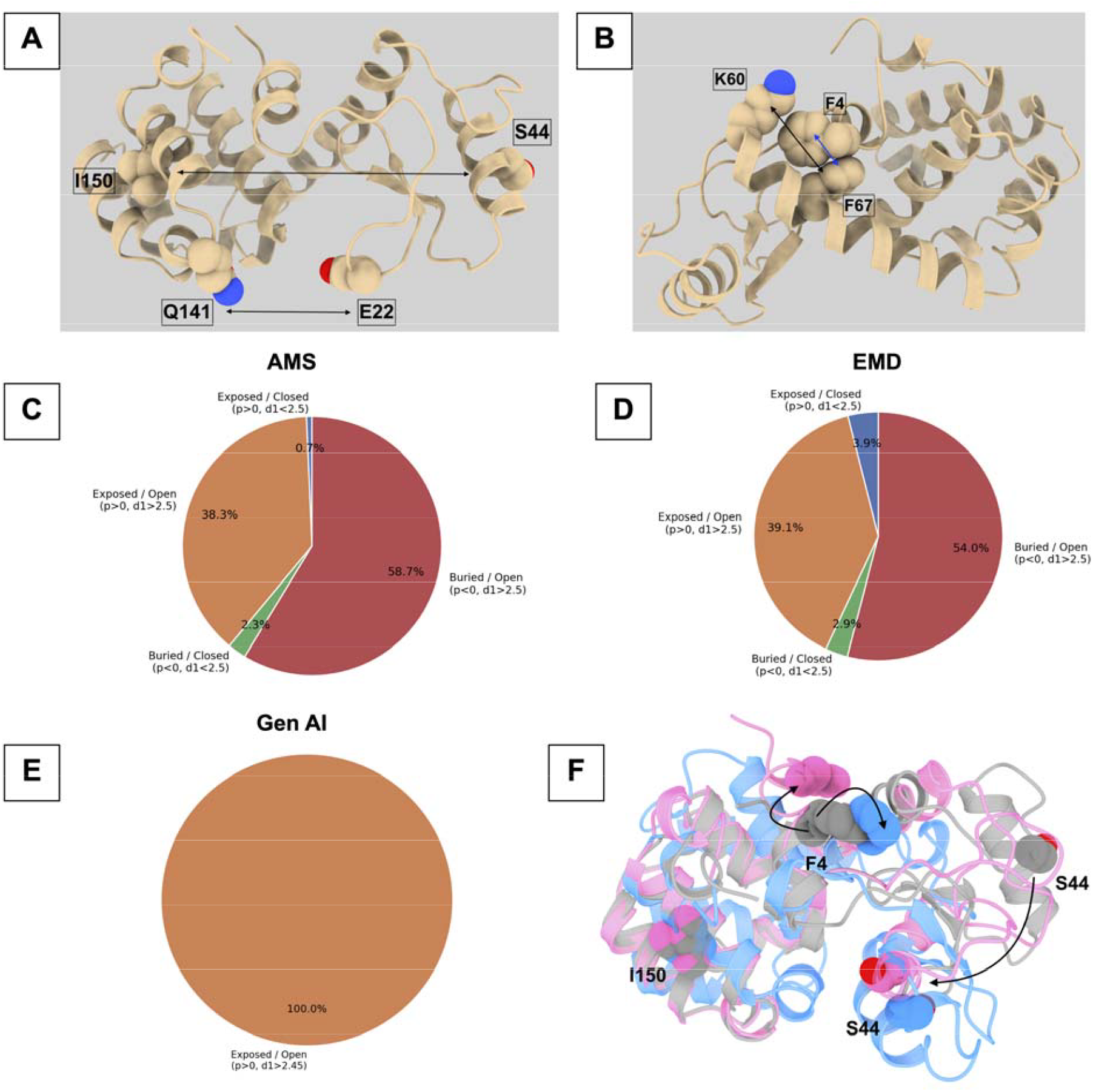
Collective variables used to characterize T4 lysozyme conformational heterogeneity and comparison of sampling across different approaches. **A:** Pictorial representation of the **d1** and **d2** collective variables, defined as the Cα–Cα distances between Ser44 (S44) and Ile150 (I150) and between Glu22 (E22) and Gln141 (Q141), respectively. The *d1* coordinate distinguishes closed and open conformations, with *d1<2*.*5*nm corresponding to the closed state and *d1>2*.*5*nm corresponding to the open state. **B:** Definition of the locking coordinate *p*, which describes the position of the Phe4 (F4) side chain relative to the hydrophobic cavity. The cavity direction is defined by the Cα atoms of Lys60 and Phe67, while the position of Phe4 is represented by the center of mass of the carbon atoms in its phenyl ring. Positive values of *p* indicate that the Phe4 side chain is solvent-exposed, whereas negative values indicate that it is buried within the hydrophobic cavity. **C–E:** Comparison of conformational sampling by AMS, EMD, and generative AI ensembles alone. AMS and EMD sample the full spectrum of conformational heterogeneity, whereas the combined generative AI ensembles predominantly sample a single state. **F:** Superposition of crystal T4 lysozyme structure shown in dark gray with representative exposed/closed and buried/closed conformations shown in magenta and blue, respectively. The comparison highlights conformational changes involving Ser44 and Phe4, which contribute to the observed conformational heterogeneity

In contrast, generative AI ensembles alone remained predominantly trapped in the exposed/open state, failing to access the transiently populated closed and buried states. Projection of generative AI ensembles onto the *d1* and *d2* CV space revealed that AF-cluster provided substantially broader conformational coverage than all other generative AI methods (**Figure 3**). This suggests that the enhanced sampling efficiency of AMS is predominantly driven by the ability of AF-cluster to populate high-energy intermediate structures that serve as productive seeds for subsequent MD simulation. To test this hypothesis directly, we repeated the AMS protocol with AF-cluster excluded from the initial ensemble. The resulting simulations failed to recover the conformational heterogeneity captured by the full AMS protocol (Figure S1, Supporting Information).

**Figure 3.**
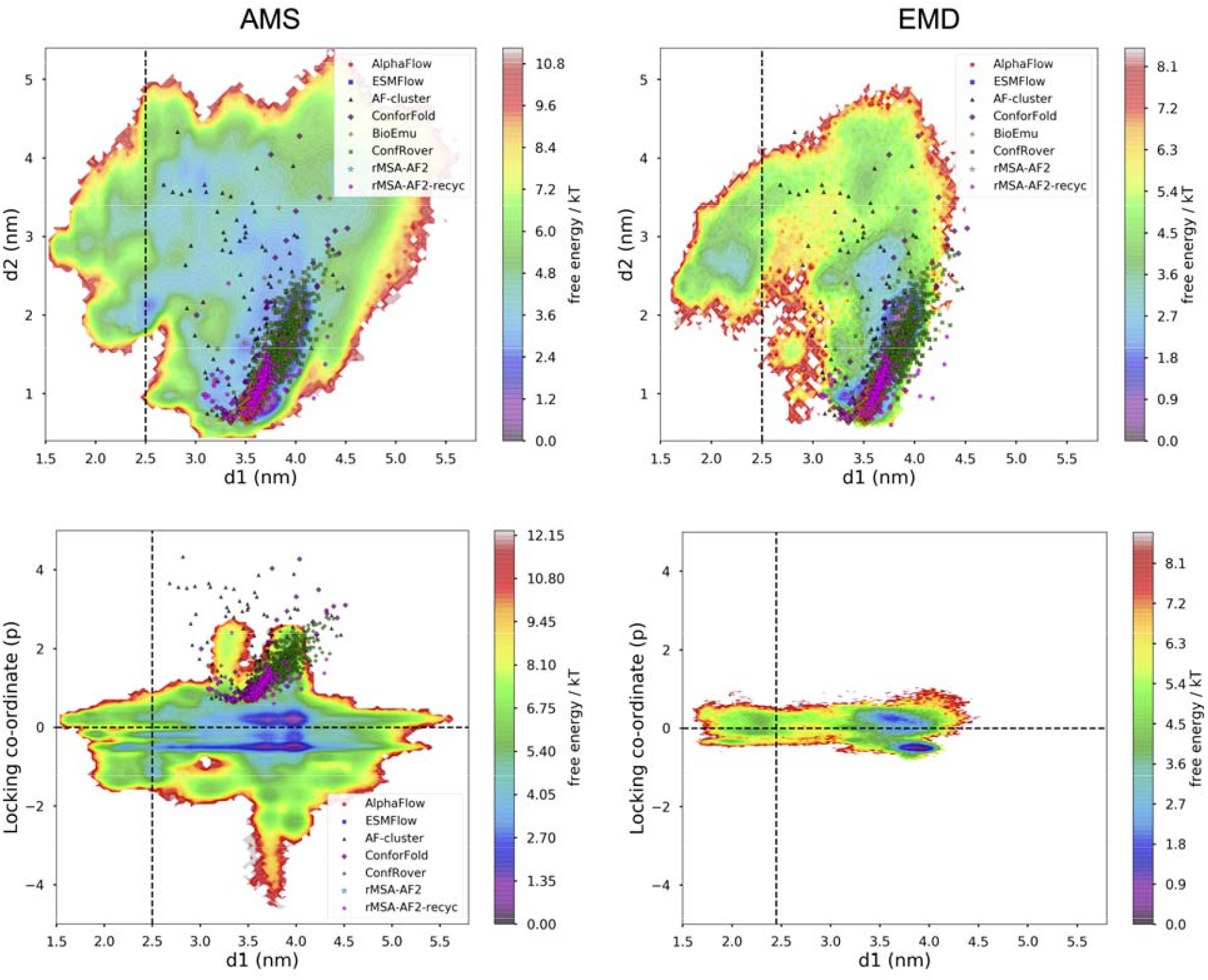
MSM-weighted free-energy landscapes projected onto different collective variables compare conformational sampling by the *AMS* and *EMD* approaches. The upper panels show equilibrium populations projected along the *d1* and *d2* collective variables, indicating that both *AMS* and *EMD* sample the closed state of T4 lysozyme, defined by *d1* < 2.5 nm. The black dotted line marks the closed–open boundary at *d1* = 2.5 nm. The lower panels show the projection of the locking coordinate *p* along *d1*, demonstrating that both *AMS* and *EMD* sample conformations with exposed and buried Phenylalanine 4 side-chain states, corresponding to *p* > 0 and *p* < 0, respectively, across both closed and open T4 lysozyme conformations. Data points from the generative AI ensembles are projected onto the physics-refined free-energy surfaces to assess the conformational coverage of each method. Among the generative AI approaches, *AF-cluster* samples a substantially broader conformational landscape than the other methods.

To further validate AMS against experiment, we computed the MSM-weighted smFRET distance distribution for the Ser44–Ile150 pair and compared it directly with experimental measurements (Figure S2 in Supporting Information). AMS recovered the transiently populated closed state in quantitative agreement with experiment. This result is significant because the EMD simulations of *Miller and co-workers(23)* were explicitly seeded with the objective of sampling the closed state using enhanced sampling, yet AMS achieved comparable recovery of this rare state starting from unbiased MD alone. The ability to reproduce an experimentally observable transiently populated conformation without prior knowledge of the target state demonstrates that AMS can access rare conformational events that are inaccessible to generative AI methods alone.

## Conclusions

This work introduces T4L as a community benchmark for evaluating methods that claim to capture conformational heterogeneity. T4L is uniquely suited to this role: its experimentally validated transient states, well characterized CVs, and extensive structural coverage in the Protein Data Bank collectively eliminate the confounding factors that have hampered fair comparisons in the field. The four-state classification framework presented here provides a transferable, physically interpretable scoring scheme that can be applied consistently across any method that generates structural ensembles.

The results presented here draw a clear boundary between what generative AI can and cannot do in its current form. Majority of the generative AI methods alone reproduce the dominant ground state (except AF-cluster which samples multiple high energy transient states) but fail to sample transiently populated closed state in T4L and their relative populations. AMS, by using generative AI ensembles as seeds rather than as end products, recovers the full conformational landscape with populations that are quantitatively comparable to those from physics-based enhanced sampling. Two rounds of unbiased MD simulation, initiated from generative AI derived starting structures, were sufficient to achieve this. This is a practically important result: AMS requires no knowledge of the target state and no enhanced sampling bias, yet it matches the performance of the EMD protocol that was explicitly designed to find the closed state. Further, the seeding strategy that combines all the conformational ensembles generated by different generative AI algorithms alleviates the limitations of each one of them and provides a much wider distribution of conformations as initial seeds, which accelerates sampling when combined with physics-based molecular simulations

Looking forward, T4L and the benchmarking framework established here are well-positioned to evaluate the next generation of hybrid methods. These include generative AI-seeded weighted ensemble simulations(24), inference-time enhanced sampling approaches such as Boltz-MetaDiffusion(25), and related methods that either combine generative AI with physics-based sampling protocols or bias the generative process during inference to access alternate states. The framework is equally applicable to MSA subsampling strategies applied to newer versions of foundation models such as AlphaFold3(26) and OpenFold3(27), where the impact of richer training data and improved architecture on conformational diversity remains an open question. As these methods mature, rigorous benchmarking against experimentally validated populations along physically meaningful CVs will be essential to distinguish genuine advances in conformational sampling from improvements in ground-state structure prediction. T4L, with its tractable size, rich experimental data, and well-defined conformational heterogeneity, provides the ideal system for that purpose.

## Methods

### Ensemble generation

Structural ensembles of T4 lysozyme (T4L) were generated from the wild-type T4L sequence using multiple generative AI-based approaches, as summarized in **Table 1**. The input amino acid sequence used for all methods was:

MNIFEMLRIDEGLRLKIYKDTEGYYTIGIGHLLTKSPSLNAAKSELDKAIGRNCNGVITKDEAEKLFNQDVDAAVRGILRNAKLKPVYDSLDAVRRCALINMVFQMGETGVAGFTNSLRMLQQKRWDEAAVNLAKSRWYNQTPNRAKRVITTFRTGTWDAYKNL

**Table 1.** Generative AI methods used for T4L ensemble generation, number of generated structures, and corresponding codebase or implementation.

| Methods | Number of structures | Codebase/implementation |
| --- | --- | --- |
| AF-cluster | 132 | https://github.com/HWaymentSteele/AF_Cluster |
| AlphaFlow | 500 | https://github.com/bjing2016/alphaflow<br>(model: AlphaFlow-MD) |
| ConforFold | 600 | https://github.com/strauchlab/ConforFold |
| ConfRover | 500 | https://github.com/ByteDance-Seed/ConfRover |
| ESMFlow | 100 | https://github.com/bjing2016/alphaflow<br>(model: ESMFlow-MD) |
| BioEmu | 499 | https://github.com/microsoft/bioemu |
| rMSA-AF2 | 80 | Colabfold (MSA: 8:16, num_recycles=6) |
| rMSA-AF2-recyc | 285 | AlphaFold2 advanced v2 (MSA: 8:16, num_recycles=3) |

The generated ensembles included both multiple-sequence-alignment-based approaches and generative models trained to produce conformationally diverse protein structures. For each method, the number of generated structures and the corresponding codebase or implementation are listed in Table 1. All structures generated across these methods were pooled into a combined generative AI ensemble for subsequent transformed-distance analysis, dimensionality reduction, and clustering.

### Definitions of collective variables

The opening and closing motion of T4 lysozyme was characterized using the Cα–Cα distance between Ser44 and Ile150, denoted as *d1*. Conformations with *d1* < 2.5 nm were classified as closed, whereas conformations with *d1*> 2.5 nm were classified as open. A second Cα–Cα distance, measured between Glu22 and Gln141 and denoted as *d2*, was used as an orthogonal projection coordinate to describe hinge-domain motion, following the collective-variable definition proposed by *Abou-Hatab and Abrams(28)*.

The solvent exposure of Phe4 was quantified using the locking coordinate, *p*. For each trajectory frame, *p* was calculated by projecting the vector from the Cα atom of Phe67 to the center of the Phe4 phenyl ring onto the vector connecting the Cα atoms of Lys60 and Phe67. The center of the Phe4 phenyl ring was defined as the geometric center of the CG, CD1, CD2, CE1, CE2, and CZ carbon atoms. The locking coordinate was computed as

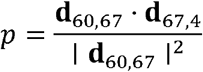

where **d**_60,67_ is the vector from the Lys60 Cα atom to the Phe67 Cα atom, and **d**_67,4_ is the vector from the Phe67 Cα atom to the center of the Phe4 phenyl ring. Positive values of *p* (*p > 0*) indicate that the Phe4 side chain is solvent-exposed, whereas negative values indicate that the side chain is buried (*p < 0)* within the hydrophobic cavity.

### Feature selection, dimensionality reduction and clustering

To construct a distance-based representation of conformational variability, structures generated by the different generative AI methods were first pooled into a single all-atom ensemble. For every structure in this combined ensemble, all Cα–Cα pairwise distances were converted into continuous contact-like variables using a smooth switching function, following *Bhakat et al(29)*.

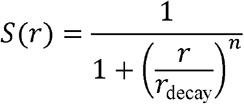

Here, *r* denotes the Cα–Cα distance, *r*_decay_ defines the distance scale over which the contact changes from formed to broken, and *n* controls the steepness of this transition. Distances much shorter than *r*_decay_ produce values close to 1, corresponding to a formed contact, whereas distances much longer than *r*_decay_ produce values close to 0, corresponding to a broken contact. This transformation replaces raw distances with a smooth and noise-tolerant measure of contact strength. We used *r*_decay_ *=* 0.90 nm and *n=* 6.

The same distance-to-contact transformation was used at two stages of the workflow. First, it was applied to the combined generative AI ensemble to identify structurally diverse starting conformations for molecular simulation. Second, after the unbiased molecular dynamics simulations were completed, the resulting trajectories were processed using the same transformed-distance representation to identify conformations for the physics-refined ensemble. Thus, both the initial generative AI predicted structures and the MD-derived conformations were analyzed in a common feature space.

For each Cα–Cα pair, the transformed contact values were evaluated across the relevant structural dataset, either the pooled generative AI ensemble or the post-processed simulation trajectories. Contacts were retained only if they showed evidence of both formation and disruption, according to the criterion:

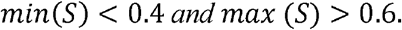

This filtering step removes distance pairs that remain essentially unchanged, either always formed or always broken, and keeps only contacts that report meaningful conformational rearrangements. In this representation, values of *S*(*r*) ≥ 0.6 correspond to formed contacts, values of *S*(*r*) ≤ 0.4 correspond to broken contacts, and intermediate values describe partially formed or fluctuating contacts.

The filtered set of transformed distance features was then used for dimensionality reduction with Slow Feature Analysis (SFA), as introduced for biomolecular simulation data by Vats et al(30). SFA identifies linear combinations of input features that change slowly over time, thereby emphasizing collective motions associated with long-timescale conformational rearrangements. For an input signal *c*(*t*), SFA generates output coordinates *y*_*k*_ (*t*) *g*_*k*_ (*c*(*t*)) by minimizing temporal variation:

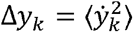

Applied to the transformed Cα–Cα contact features, SFA identified the slowest collective distance patterns that distinguish conformationally heterogeneous regions of T4 lysozyme. The first two slow features were used as the reduced feature space for K-center clustering(31). Cluster centers selected from the combined generative AI ensemble were saved as PDB structures and used as starting points for the first round of molecular dynamics simulations. The same procedure was subsequently applied during post-processing of the simulation trajectories to select the physics-refined ensemble for the next round of simulations. SFA and K-center clustering were performed using the MDML software package (GitHub: https://github.com/svats73/mdml/tree/main)

### Molecular dynamics simulations

Each selected structure from starting ensemble or physics-refined ensemble was prepared using the *tleap* module in *Amber2022(32, 33)*, following the general simulation protocol described by *Meller et al(34)*. Protein atoms were parameterized with the *AMBER ff14SB(35)* force field. Each system was neutralized with the appropriate counterions and solvated in a truncated-octahedron box of *TIP3P* water molecules, with a minimum distance of 10 Å between any protein atom and the edge of the simulation box.

Energy minimization was carried out in two steps. First, solvent molecules and ions were minimized while harmonic restraints were applied to the protein atoms using a force constant of 100 kcal mol□^1^ Å□^2^. This was followed by an unrestrained minimization of the full solvated system. The resulting *Amber* topology and coordinate files were then converted to *GROMACS* format using *Acpype(36)*, and all subsequent equilibration and production simulations were performed using *GROMACS 2022(37)*.

Systems were gradually heated from 0 to 300 K over 500 ps in the NVT ensemble while applying positional restraints to backbone heavy atoms with a force constant of 500 kJ mol□^1^ nm□^2^. After heating, each system was equilibrated for 200 ps in the NPT ensemble at 300 K and 1 bar without positional restraints. Temperature was maintained using the velocity-rescale thermostat, and pressure was controlled using the *Parrinello–Rahman* barostat(38).

Unbiased production simulations were then performed in the NPT ensemble using a 2 fs integration time step. Nonbonded interactions were treated with a 1.0 nm cutoff, long-range electrostatics were calculated using the particle-mesh Ewald method(39), and bonds involving hydrogen atoms were constrained using the LINCS(40) algorithm. Production trajectories were initiated from the SFA selected cluster centers and ran for 200 ns each, with coordinates saved every 10 ps.

### Enhanced Molecular Dynamics simulations

The molecular dynamics simulations used in this study were adapted from *Miller et al(23)*. Briefly, the dataset comprises a 5 μs unbiased MD trajectory, which did not sample the open-to-closed conformational transition. To drive the transition, the authors applied a metadynamics bias along the Cα–Cα distance between Ser44 and Ile150, extracted snapshots along the resulting free-energy pathway, and launched short unbiased MD simulations from each. For our analysis, we did not use the metadynamics trajectories themselves; instead, we combined the 5 μs unbiased trajectory with the short unbiased simulations seeded along the biased pathway.

### Markov state model

Markov state models (MSMs) were constructed using the PyEMMA(41) package with RMSD-based featurization of both AMS and EMD trajectories. K-means clustering (*K = 200*) was employed to discretize conformational space, and state populations were computed across multiple lag times to assess convergence and compare the relative equilibrium distributions of AMS and EMD ensembles (Figure S3 and S4 in Supporting Information).

### smFRET prediction

smFRET prediction was carried out using Enspara(42) software package (https://enspara.readthedocs.io/en/latest/smFRET.html). Trajectories from AMS were featurized by transformed distances followed by SFA (as described in “*Feature selection, dimensionality reduction and clustering*” subsection) followed by K-means clustering with k=500. smFRET was predicted using the protocol described by *Miller et al(23)*.

## Competing interests

Authors declares no conflict of interests.

## Acknowledgements

The author thanks Justin J. Miller of the University of Pennsylvania for performing the smFRET calculations and providing the dataset corresponding to the EMD simulations. Further acknowledgement goes to Prof. Eva M. Strauch and Dr. Raulia Syrlybaeva of Washington University in St. Louis for providing the ConforFold ensemble, and to Shray Vats of Boston University for providing the ConfRover ensemble.

## Data availability

Generative AI ensembles, closed states of T4L and codes to calculate the distance, RMSD and locking co-ordinate can be accessed here: https://doi.org/10.5281/zenodo.20111229

## Notes

https://doi.org/10.5281/zenodo.20111229

### Competing Interest Statement

The authors have declared no competing interest.

