## Supporting Information for "Benchmarking generative AI and physics based molecular simulation for sampling conformational heterogeneity in T4 Lysozyme"

Soumendranath Bhakat

AlloTec Bio Inc., USA

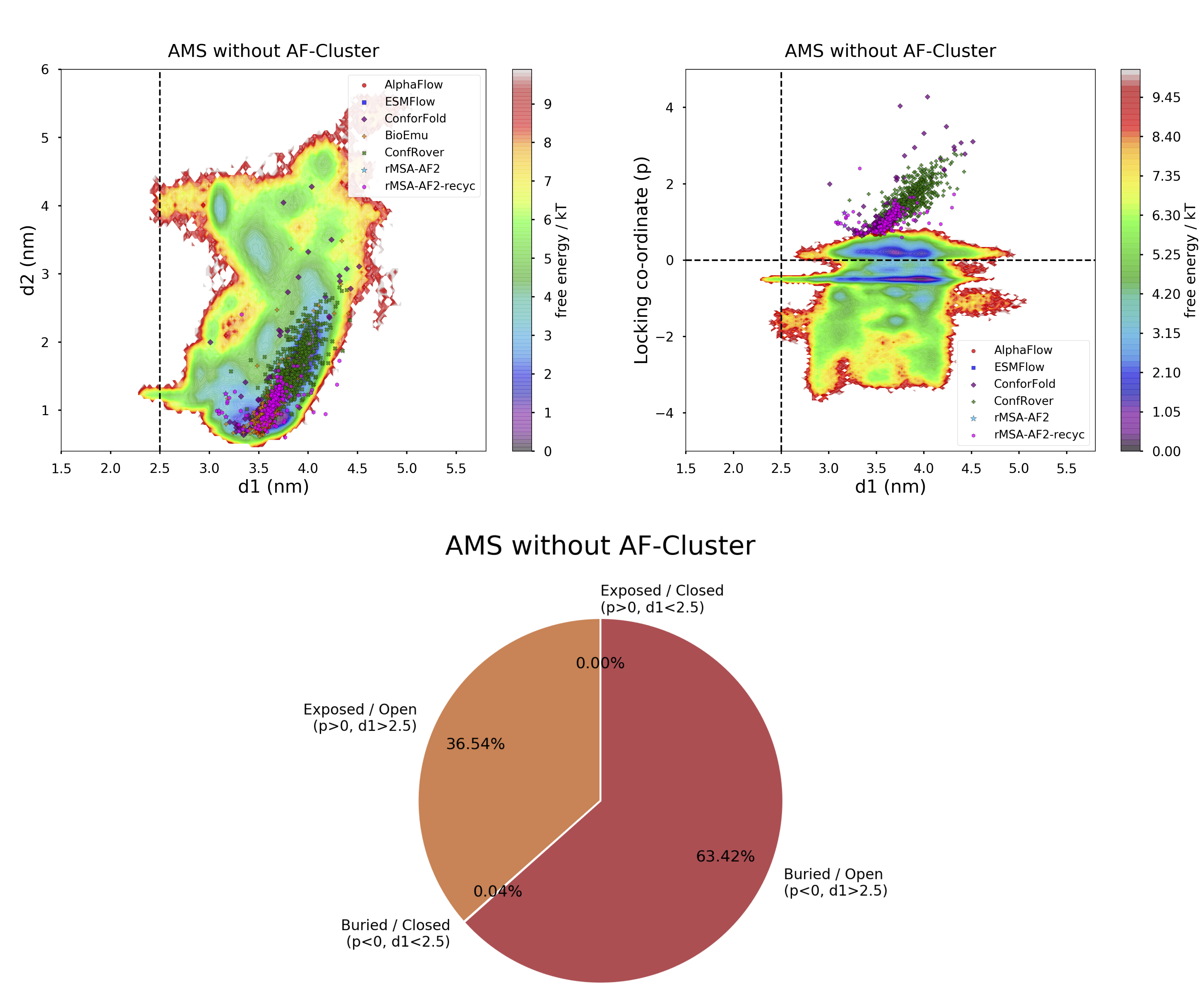

**Figure S1**. MSM-weighted free energy surfaces projected along different CVs show that molecular simulations initiated from the generative AI ensemble without AF-cluster fail to sample the closed state, in contrast to AMS simulations that include AF-cluster (see Figures 2 and 3 in the main text).

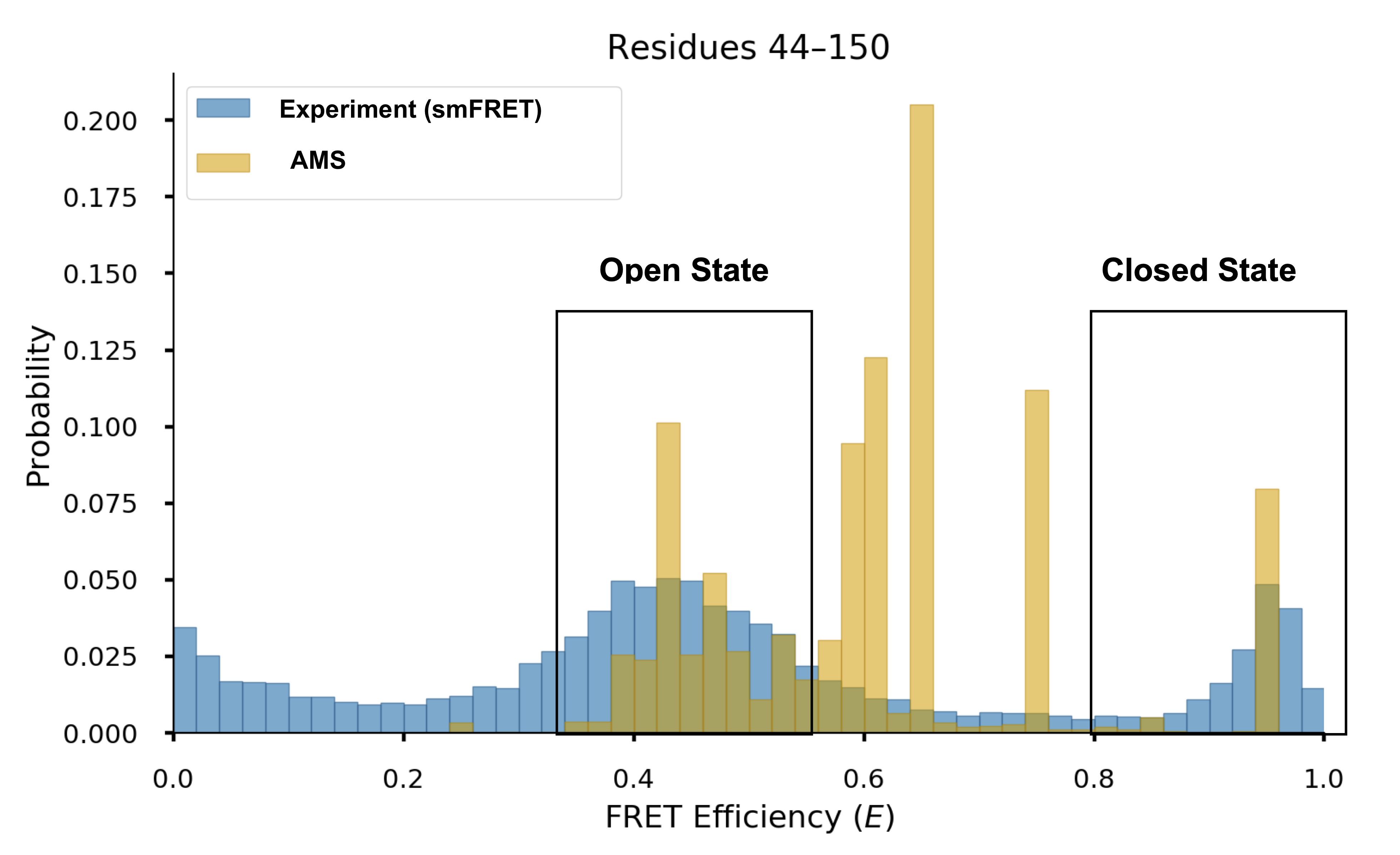

Figure S2. Comparison between MSM-weighted FRET efficiency from AMS and experimental smFRET shows sampling of both open and closed states in T4L. Notably, AMS captured the smFRET probability corresponding to the closed state, a rare event previously sampled only by EMD molecular simulations reported by *Bowman et al*.

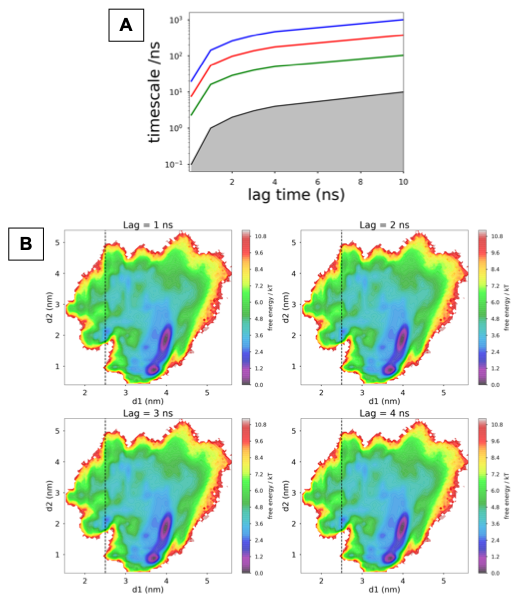

Figure S3. Lag time vs timescale plot and projection of MSM weighted free energy surface across different lag times highlighting convergence of AMS.

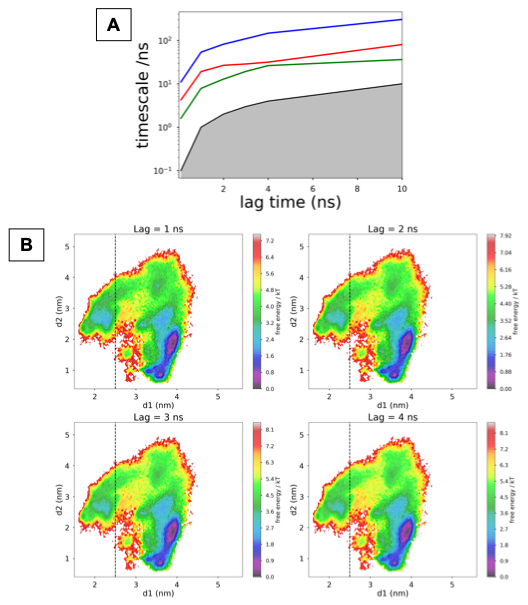

Figure S4. Lag time vs timescale plot and projection of MSM weighted free energy surface across different lag times highlighting convergence of EMD simulations.
